# Beyond differences in means: robust graphical methods to compare two groups in neuroscience

**DOI:** 10.1101/121079

**Authors:** Guillaume A. Rousselet, Cyril R. Pernet, Rand R. Wilcox

## Abstract

If many changes are necessary to improve the quality of neuroscience research, one relatively simple step could have great pay-offs: to promote the adoption of detailed graphical methods, combined with robust inferential statistics. Here we illustrate how such methods can lead to a much more detailed understanding of group differences than bar graphs and t-tests on means. To complement the neuroscientist’s toolbox, we present two powerful tools that can help us understand how groups of observations differ: the shift function and the difference asymmetry function. These tools can be combined with detailed visualisations to provide complementary perspectives about the data. We provide implementations in R and Matlab of the graphical tools, and all the examples in the article can be reproduced using R scripts.

## Introduction

Despite the potentially large complexity of experiments in neuroscience, from molecules, neurones, to large scale brain measurements and behaviour, data pre-processing and subsequent analyses typically lead to massive dimensionality reduction. For instance, reaction time distributions are summarised by their means, so they can be compared easily across conditions and participants; the firing rate of individual neurones is averaged in a time-window of interest; BOLD signal is averaged in a region of interest. Because of such complexity reduction, researchers often focus on a limited number of group comparisons, such that the thrust of an article tends to depend on a few distributions of continuous variables. In addition, our own experience, as well as surveys of the literature (Allen *et al.*, 2012; Weissgerber *et al.*, 2015), suggest data representation standards need an overhaul: the norm is to hide distributions behind bar graphs, using the standard deviation or the standard error of the mean to illustrate uncertainty. That standard, coupled with the dominant use of t-tests and ANOVAs on means, can mask potentially rich patterns. As a result, many neuroscience datasets are under-exploited.

To make the most of neuroscience datasets, we believe one solution is to adopt robust and detailed graphical methods, which could have great pay-offs for the field (Rousselet *et al.*, 2016b). Briefly, modern statistical methods offer the opportunity to get a deeper, more accurate and more nuanced understanding of data (Wilcox, 2017). For instance, in Figure 1, the classic combination of a bargraph and a t-test suggests the two groups of participants differ very little in cerebellum local grey matter volume (Voxel Based Morphometric data from Pernet *et al.*, 2009a). Using a more detailed graphical description such as a dotplot hints at a more interesting bimodal distribution in the patient group, and alternative analyses suggest that individual differences in patients’ grey matter volumes are related to behavioural variables (see details in Pernet *et al.*, 2009b). In the rest of the article we cover other examples in which alternative methods are more informative than t-tests. In addition, even when t-tests are appropriate for the problem at hand, they lack robustness, as illustrated in this simple example. Imagine we have a vector of observations [1, 1.5, 1.6, 1.8, 2, 2.2, 2.4, 2.7] and null hypothesis of 1. The one-sample t-test on mean gives t = 4.69, p= 0.002 and 95% confidence interval = [1.45, 2.35]. A single outlier can have devastating effects: for instance, adding the observation 8 to our previous vector now leads to t = 2.26, p = 0.054, and 95% confidence interval = [0.97, 4.19]. In this latter case, we fail to reject, despite growing evidence that we are not sampling from a distribution with mean of 1. Yet, inferential tools robust to outliers and other distribution problems are readily available and have been described in many publications (Wilcox & Keselman, 2003; Erceg-Hurn & Mirosevich, 2008; Wilcox, 2009). The examples above also illustrate why detailed descriptions of distributions can be vital to make sense of a dataset, without relying blindly on a unique inferential test, which might be asking the wrong question about the nature of the effects.

**Figure 1.**
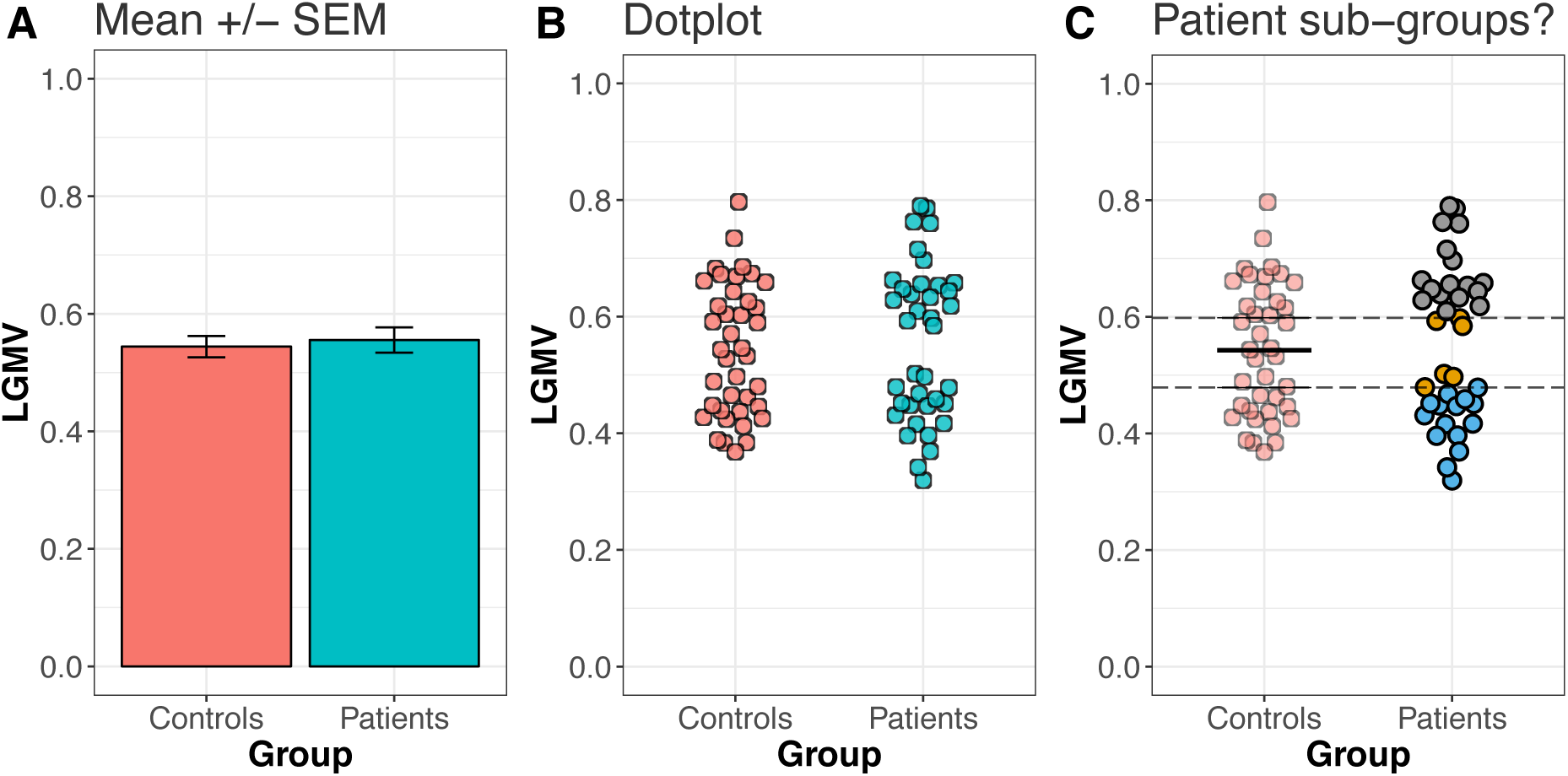
Beyond bar graphs and t-tests. Data from Pernet et al. (2009a), showing the local grey matter volume (LGMV) in the cerebellum of control participants and of participants with dyslexia (patients). **A. Bargraph and t-test suggest that the two groups do not differ**: *t* = −0.4, df = 72.6, p-value = 0.692, difference = −0.01 [−0.07, 0.04]. **B. A dotplot suggests a bimodal distribution in patients.** Each point is a participant and the points were jittered to reduce overlap. A dotplot is also called a stripchart or a 1 dimensional scatterplot. **C. An alternative analysis suggests sub-groups of patients.** Using the controls as reference, we can sort patients into subgroups, based on whether they fall above (grey), within (orange), or below (blue) certain limits. For instance, here we used the confidence interval of the median of the control group as a reference to classify the patients. Using the mean instead of the median, all patients would fall outside the control confidence interval, as reported in Pernet et al. (Pernet *et al.*, 2009b).

The benefits of illustrating data distributions have been emphasised in many publications and is often the topic of one of the first chapters of introductory statistics books (Wilcox, 2006; Allen *et al.*, 2012; Duke *et al.*, 2015; Weissgerber *et al.*, 2015; Cook *et al.*, 2016). One of the most striking examples is provided by Anscombe’s quartet (Anscombe, 1973), in which very different distributions, illustrated using scatterplots, are associated with the same summary statistics. The point of Anscombe’s quartet is simple and powerful, yet often underestimated: unless results are illustrated in sufficient details, standard summary statistics can lead to unwarranted conclusions.

As demonstrated by the Anscombe’s quartet, it is easy to fool ourselves if we use the wrong tools, because they ask the wrong questions. Take for instance Figure 2, which illustrates a few examples of how distributions can differ. Obviously, distributions can differ in other aspects than those illustrated, and in combinations of these aspects, as we will explore in other examples in the rest of this article. Yet, despite these various potential patterns of differences, the standard group comparison using t-test on means makes the very strong assumptions that the most important difference between two distributions is a difference in central tendency, and that this difference is best captured by the mean. This is clearly not the case if distributions differ in spread or skewness, as illustrated in the caricatural examples of columns 3 and 4 of Figure 2.

**Figure 2.**
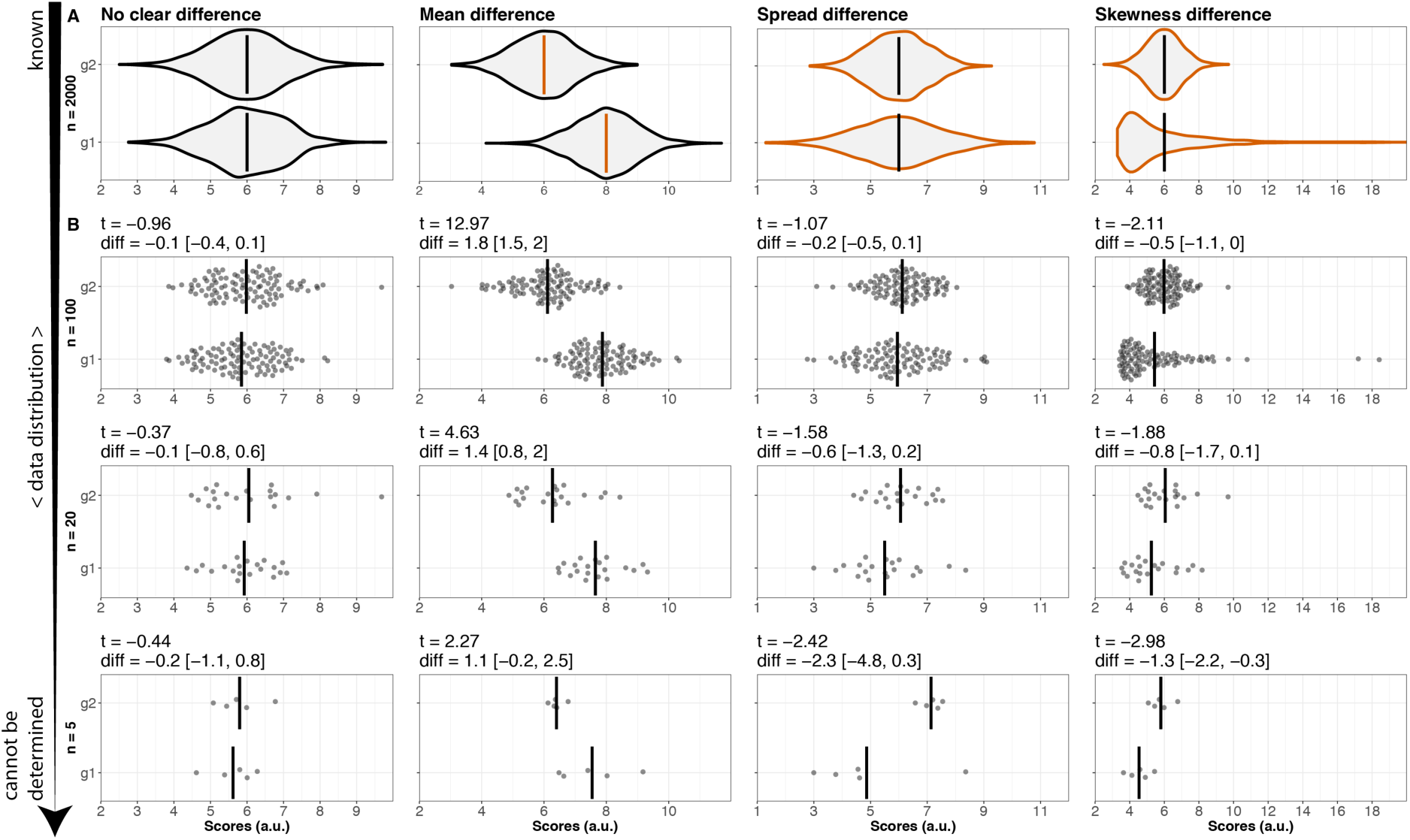
Distribution differences and sample sizes. A. Distributions can differ in other aspects than the mean. Columns show distributions that differ in four different ways. Each example portrays two randomly generated populations, each with n = 2000. In examples 1, 3 and 4, the two distributions have the same mean. In example 2, the means of the two distributions differ by 2 arbitrary units. In examples 3 and 4, the distributions differ in shape. The distributions are illustrated with violinplots. The vertical bars indicate the mean of each distribution. Orange indicates differences in mean or in shape. **B. Data distributions cannot be estimated with very small sample sizes.** The three rows illustrate random subsamples of data from panel A, with sample sizes n = 100, n = 20, and n = 5. Above each plot, the t value, mean difference and its confidence interval are reported. The vertical bars indicate the mean of each sample. On the left of the figure, the downward pointing arrow illustrates the decreasing certainty about the shape of the distribution.

The problem with asking a very narrow question about the data using a t-test on mean is exacerbated by the small sample sizes common in neuroscience. Small sample sizes are associated with low statistical power, inflated false discovery rate, inflated effect size estimation, and low reproducibility (Button *et al.*, 2013; Colquhoun, 2014; Forstmeier *et al.*, 2016; Munafò *et al.*, 2017; Poldrack *et al.*, 2017). Small sample size also prevents us from properly estimating and modelling the populations we sample from. Consequently, small n stops us from answering a fundamental, yet often ignored empirical question: how do distributions differ?

Let’s consider the n = 2000 populations in Figure 2A. If we draw random sub-samples of different sizes from these populations (Figure 2B), we can get a sense of the sorts of problems we might be facing as experimenters, when we draw one sample to try to make inferences about an unknown population. For instance, even with 100 observations we might struggle to approximate the shape of the parent population. Without additional information, it can be difficult to determine if an observation is an outlier, particularly for skewed distributions. And in column 4 of Figure 2, the samples with n = 20 and n = 5 are very misleading. Nevertheless, some of the techniques described below can be applied to sample sizes as low as 10 or 20 observations – see section Recommendations for details.

All the figures in this article are licensed CC-BY 4.0 and can be reproduced using scripts in the R programming language (R Core Team, 2016) and datasets available on figshare (Rousselet *et al.*, 2016a). The figshare repository also includes Matlab code implementing the main R functions. The main R packages used to make the figures are *ggplot2* (Wickham, 2016), *cowplot* (Wilke, 2016), *ggbeeswarm* (Clarke & Sherrill-Mix, 2016), *retimes* (Massidda, 2013), and *rogme* (Rousselet & Wilcox, 2016), which was developed for this article.

## Beyond the mean: a matter of perspectives

The previous examples illustrate that to understand how distributions differ, large sample sizes are needed. How large is partly an empirical question that should be addressed in each field for different types of variables. We will make a few recommendations at the end of this article. For now, assuming that we have large enough sample sizes, why do we need to look beyond the mean? And how do we go about quantifying how distributions differ? It’s a matter of perspectives.

When comparing two independent groups, we can consider different perspectives; yet one tends to dominate, as we typically ask:

> ‘How does the typical observation/participant in one group compares to the typical observation/participant in the other group?’ *(Question 1)*.

To answer this question, we compare the marginal distributions using a proxy: the mean. Indeed, following this approach, we simply summarise each distribution by one value, which we think provides a good representation of the average Joe in each distribution. An interesting alternative approach consists in asking:

> ‘What is the typical difference (effect) between any member of group 1 and any member of group 2?’ *(Question 2)*.

In other words, if we randomly select one member of group 1 and one member of group 2, by how much do they differ? This comparison can be done by systematically comparing members of the two groups and summarising the distribution of pairwise differences by using one value, for instance the mean. This perspective is particularly useful in a clinical setting, to get a sense of how a randomly selected patient tends to differ from a randomly selected control participant; or to compare young vs. old rats for example.

To answer *Question 1* or *Question 2*, it is essential to appreciate that there is nothing special about using the mean to summarise distributions. The mean is one of several options for the job, and often not the best choice. Indeed, the mean is not robust to outliers, and robust alternatives such as medians, trimmed means and M-estimators are more appropriate in many situations (Wilcox, 2017). Similarly, the standard least squares technique underlying t-tests and ANOVAs is often inappropriate because its assumptions are easily violated (Wilcox, 2001; Erceg-Hurn & Mirosevich, 2008). Also, there is no reason to limit our questioning of the data to the average Joe in each distribution: we have tools to go beyond differences in central tendency, for instance to explore effects in the tails of the distributions. We can thus ask a more detailed version of *Question 1*: ‘How do observations in specific parts of a distribution compare between groups?’. We can tackle this more specific question by performing systematic group comparisons using a shift function, a tool that we will present in detail in the next section. *Question 2* can also be extended by quantifying multiple aspects of the distribution of differences, including its symmetry, which can be assessed using the difference asymmetry function, introduced later in this article.

We can ask similar questions for dependent groups. Dependent groups could involve the same participants/animals tested in two experimental conditions, or in the same condition but at different time points, for instance before and after an intervention. When considering dependent groups, two main questions are usually addressed:

> ‘How does the typical observation in condition 1 compare to the typical observation in condition 2?’ *(Question 1)*.
>
> ‘What is the typical difference (effect) for a randomly sampled participant?’ (*Question 2)*.

Interestingly these two questions lead to the same answers if the mean is used as a measure of central tendency: the difference of two means is the same as the mean of the differences. That’s why a paired t-test is the same as a one-sample test on the pairwise differences. However, if other estimators are used, or other aspects of the distributions are considered, the answers to the two questions can differ. For instance, the difference between the medians of the marginal distributions is usually not the same as the median of the differences. Similarly, exploring entire distributions can reveal strong effects not or poorly captured by the mean.

To address these different perspectives on independent and dependent groups, and to quantify how distributions differ, we propose an approach that combines two important steps. The first step is to provide more comprehensive data visualisation, to guide analyses, but also to better describe how distributions differ (Wilcox, 2006; Allen *et al.*, 2012; Weissgerber *et al.*, 2015). The second step is to focus on robust estimators and alternative techniques to build confidence intervals (Wilcox, 2017). Robust estimators perform well with data drawn from a wide range of probability distributions. This framework is focused on quantifying how and by how much distributions differ, to go beyond the binary descriptions of effects as being significant or non-significant.

## The shift function

A systematic way to characterise how two independent distributions differ was originally proposed by Kjell Doksum: to plot the difference between the quantiles of two distributions as a function of the quantiles of one group (Doksum, 1974; Doksum & Sievers, 1976; Doksum, 1977). This technique is called a shift function, and is both a graphical and an inferential method. Quantiles are particularly well-suited to understand how distributions differ because they are informative, robust and intuitive.

In 1995, Wilcox proposed an alternative technique which has better probability coverage and more statistical power than Doksum & Sievers’ 1976 approach (Wilcox, 1995). In short, Wilcox’s technique:

- uses the Harrell-Davis quantile estimator to estimate the deciles of two distributions (Harrell & Davis, 1982);
- computes 95% confidence intervals of the decile differences with a bootstrap estimation of the deciles’ standard error;
- controls for multiple comparisons so that the type I error rate remains around 5% across the nine confidence intervals (this means that the confidence intervals are larger than what they would be if the two distributions were compared at only one decile).

Figure 3 illustrates a shift function and how it relates to the marginal distributions. It shows an extreme example, in which two distributions differ in spread, not in the location of the bulk of the observations. In that case, any test of central tendency will fail to reject (e.g. one-sample t-test on means: *t* = 0.91, *p* = 0.36), but it would be wrong to conclude that the two distributions do not differ. In fact, a Kolmogorov-Smirnov test reveals a significant effect (test statistics = 0.109, critical value = 0.0607), and several robust measures of effect size would also suggest non-trivial effects (Wilcox & Muska, 2010; Ince *et al.*, 2016). This shows that if we do not know how two independent distributions differ, the default test should not be a t-test but a Kolmogorov-Smirnov test. But a significant Kolmogorov-Smirnov test only suggests that two independent distributions differ, it does not tell us how they differ.

**Figure 3.**
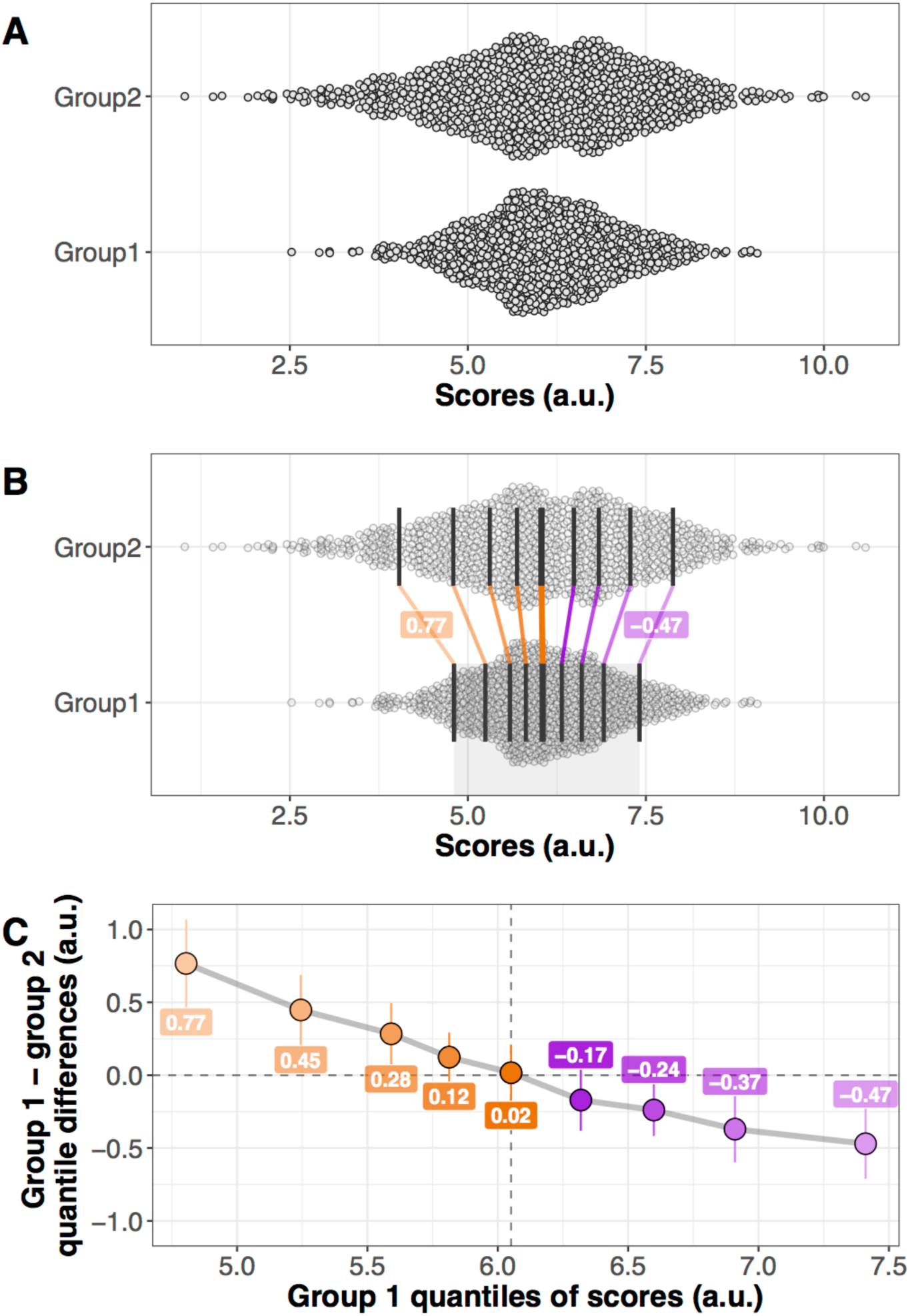
Simulated example of a pair of independent distributions and their associated shift function. **A.** Marginal distributions. The two marginal distributions (n = 1000 each) differ in spread and are illustrated using jittered 1D scatterplots (also called stripcharts or dotplots). The spread of the points is proportional to the local density of observations. The observations from each group are hypothetical scores in arbitrary units (a.u.). **B.** Same data as in panel A, but with vertical lines marking the deciles for each group. The thicker vertical line in each distribution is the median. Because of the difference in spread, the first decile of group 2 is lower than that of group 1; similarly, the ninth decile of group 2 is higher than that of group 1. Between distributions, the matching deciles are joined by coloured lined. If the decile difference between group 1 and group 2 is positive, the line is orange; if it is negative, the line is purple. The values of the differences for deciles 1 and 9 are indicated in the superimposed labels. **C.** Shift function. Panel C focuses on the portion of the x-axis marked by the grey shaded area at the bottom of panel B. It shows the deciles of group 1 on the x-axis – the same values that are shown for group 1 in panel B. The y-axis shows the differences between deciles: the difference is large and positive for decile 1; it then progressively decreases to reach almost zero for decile 5 (the median); it becomes progressively more negative for higher deciles. Thus, for each decile the shift function illustrates by how much one distribution needs to be shifted to match another one. In our example, we illustrate by how much we need to shift deciles from group 2 to match deciles from group 1. For each decile difference, the vertical line indicates its 95% bootstrap confidence interval. When a confidence interval does not include zero, the difference is considered significant in a frequentist sense, with an alpha threshold of 0.05.

The shift function can help us understand and quantify how two distributions differ. Concretely, the shift function describes how one distribution should be re-arranged to match another one: it estimates how and by how much one distribution must be shifted. In Figure 3C, the shift function shows the decile differences between group 1 and group 2, as a function of group 1 deciles. The first decile of group 1 is slightly under 5, which can be read in the top section of panel B, and on the x-axis of the shift function. The first decile of group 2 is lower; as a result, the first decile difference between group 1 and group 2 is positive: thus, to match the first deciles of the two distributions, the first decile of group 2 needs to be shifted up. Deciles 2, 3 and 4 show the same pattern, but with progressively weaker effect sizes. Decile 5 is well centred, suggesting that the two distributions do not differ in central tendency. As we move away from the median, we observe progressively larger negative differences, indicating that to match the right tails of the two distributions, group 2 needs to be shifted to the left, towards smaller values - hence the negative sign. Across quantile differences, the negative slope indicates that the two distributions differ in spread, and the steepness of the slope relates to the strength of the difference in spread between distributions. In other cases, non-linear trends would suggest differences in skewness or higher-order moments too.

To get a good understanding of the shift function, Figure 4 illustrates its behaviour in the other situations portrayed in Figure 2: no clear difference, mean difference, skewness difference. The first column of Figure 4 shows two large samples drawn from a standard normal population. In that case, a t-test on means is not significant (t=−0.45, p = 0.65), and as expected, the shift function shows no significant differences for any of the deciles. The shift function is not perfectly flat, as expected from random sampling of a limited sample size. The samples are both n = 1000, so for smaller samples even more uneven shift functions can be expected by chance. Also, the lack of significant differences should not be used to conclude that we have evidence for the lack of effect.

**Figure 4.**
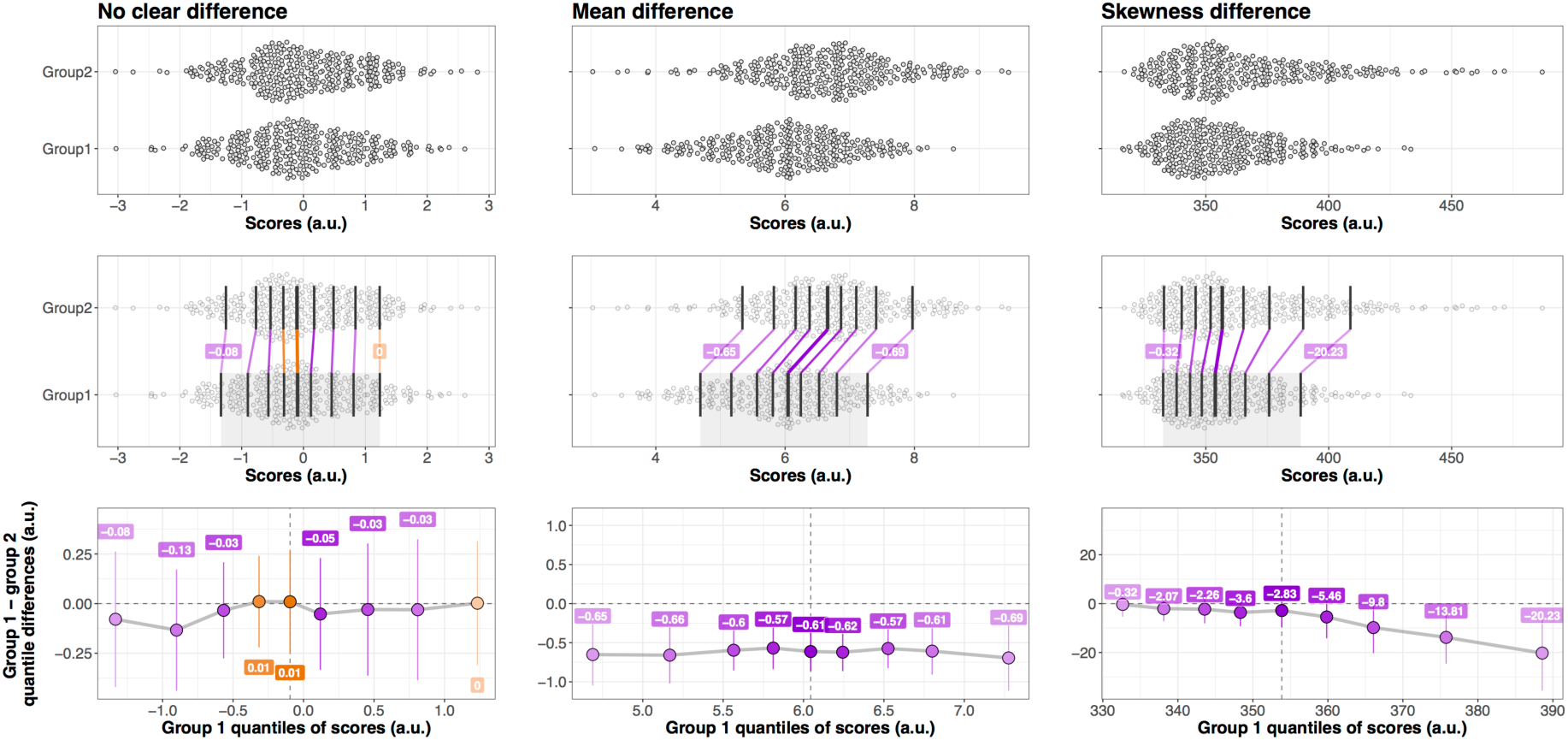
Examples of pairs of independent distributions and their associated shift functions. See details in Figure 3 caption.

In the middle column of Figure 4, the two distributions differ in central tendency: in that case, a t-test on means is significant (t=−7.56, p < 0.0001), but this is not the full story. The shift function shows that all the differences between deciles are negative and around −0.6. That all the deciles show an effect in the same direction is the hallmark of a completely effective method or experimental intervention. This consistent shift can also be described as first order stochastic ordering, in which one distribution stochastically dominates another (Speckman *et al.*, 2008). Thus, the shift function relates to the delta plot, which is an extension of Q-Q plots for the comparison of two distributions on a quantile scale (De Jong *et al.*, 1994; Ridderinkhof *et al.*, 2005; Speckman *et al.*, 2008). The shift function is also related to relative distribution methods (Handcock & Morris, 1998).

For the data presented in the third column of Figure 4, a t-test on means is significant (t=−3.74, p-value = 0.0002). However, the way the two distributions differ is very different from our previous example: the first five deciles are near zero and follow almost a horizontal line, and from deciles 5 to 9 differences increase linearly. Based on the confidence intervals, only the right tails of the two distributions seem to differ, which is captured by significant differences for deciles 8 and 9. The non-linearity in the shift function reflects these asymmetric differences.

## Neuroscience applications

### Exploration of effects

We can put the shift function in context by looking at the original example discussed by Doksum (Doksum, 1974; Doksum, 1977), concerning the survival time in days of 107 control guinea pigs and 61 guinea pigs treated with a heavy dose of tubercle bacilli (Bjerkedal, 1960) (Figure 5A). Relative to controls, the animals that died the earliest tended to live longer in the treatment group, suggesting that the treatment was beneficial to the weaker animals (decile 1). However, the treatment was harmful to animals with control survival times larger than about 200 days (deciles 4-9). Thus, this is a case where the treatment has very different effects on different animals. As noted by Doksum, the same experiment was performed 4 times, each time giving similar results. An important point, because of the increased resolution afforded by shift functions, replications are necessary to confirm specific patterns observed in exploratory work (Wagenmakers *et al.*, 2012).

**Figure 5.**
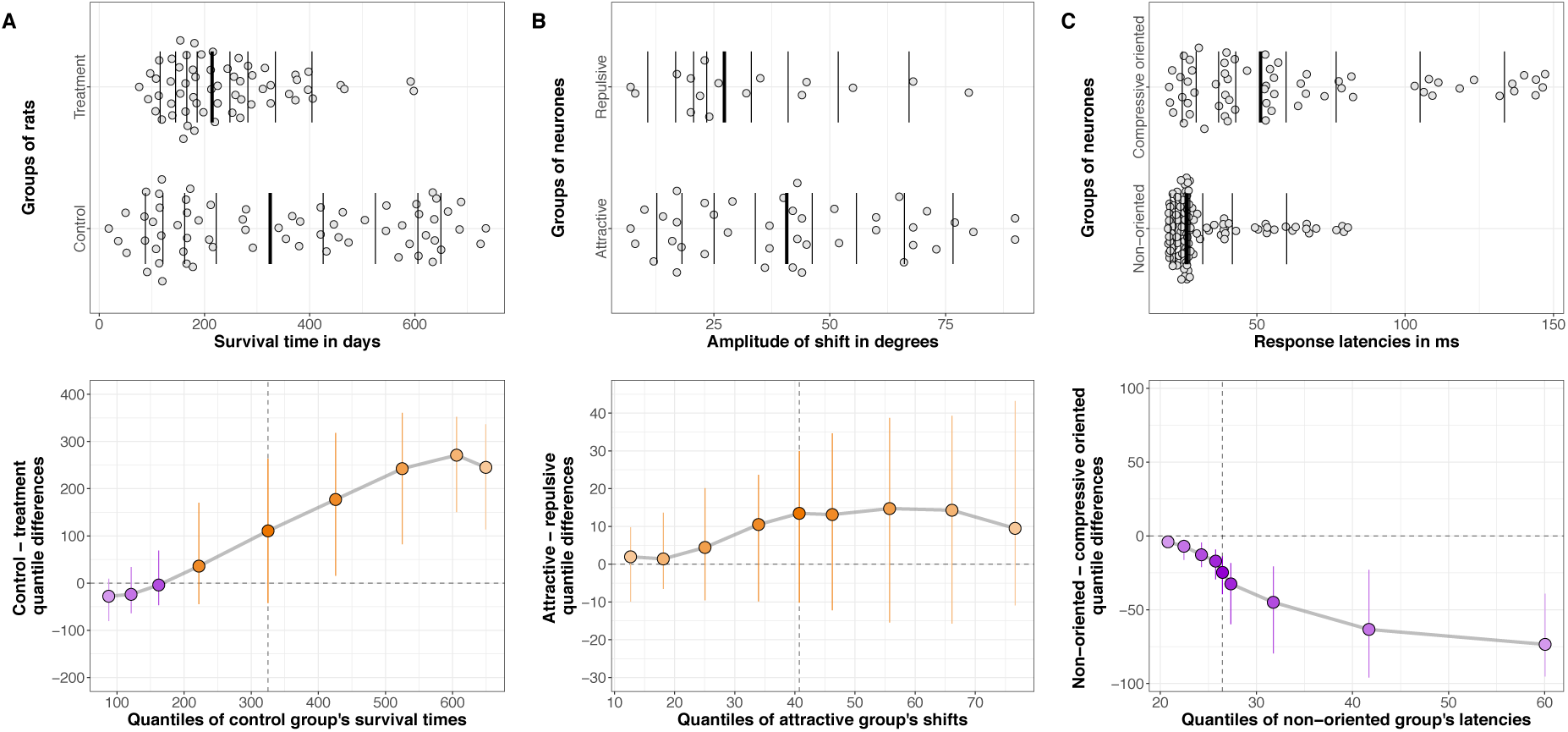
Examples of shift function applications. **A.** Data from (Bjerkedal, 1960), and used to illustrate the shift function in (Doksum, 1974). **B.** Data from Figure 5A of (Chanauria *et al.*, 2016). **C.** Data from Figure 9A of (Talebi & Baker, 2016). Data in panel A were obtained from a table in the original publication. Data from panels B and C were kindly provided by the authors. In row 1, stripcharts were jittered to avoid overlapping points. The vertical lines mark the deciles, with a thicker line for the median. Row 2 shows the matching shift functions. See other details in Figure 3 caption.

Panels B and C of Figure 5 show other examples of asymmetric effects in skewed distributions. Both panels show results from recordings from the cat visual cortex from two research groups (Chanauria *et al.*, 2016; Talebi & Baker, 2016). Panel B illustrates the adaptation response (amplitude of shift) of two independent groups of neurones with opposite responses (attractive vs. repulsive adaptation). A two-sample t-test on means is not significant (t = 1.46, p = 0.15). A shift function suggests that the two groups differ, with increasing differences for progressively larger amplitudes of shift; however, uncertainty is large. The problem would be worth exploring with a larger sample, to determine if the largest attractive shifts tend to be larger than the largest repulsive shifts. Another example of recording from the cat visual cortex is provided in panel C, in which the response latencies of two independent groups of neurones clearly differ, with much earlier latencies in non-oriented compared to compressive oriented cells. A shift function suggests a more detailed pattern: the two groups differ very little for short latencies, and progressively and non-linearly more as we move to their right tails.

Figure 5 examples are particularly important because we anticipate that, as researchers progressively abandon bar graphs for more informative alternatives (Weissgerber *et al.*, 2015; Rousselet *et al.*, 2016b), such skewed distributions and non-uniform differences will appear to be more common in neuroscience.

### Hypothesis testing

The shift function is also well suited to investigate how reaction time distributions differ between experimental interventions, such as tasks or pharmaceutical treatments. This approach requires building shift functions in every participant. Results could then be summarised, for instance, by reporting the number of participants showing specific patterns, and by averaging the individual shift functions across participants. One could imagine different situations, as illustrated in Figure 6, in which a manipulation:

- affects most strongly slow behavioural responses, but with limited effects on fast responses;
- affects all responses, fast and slow, similarly;
- has stronger effects on fast responses, and weaker ones for slow responses.

**Figure 6.**
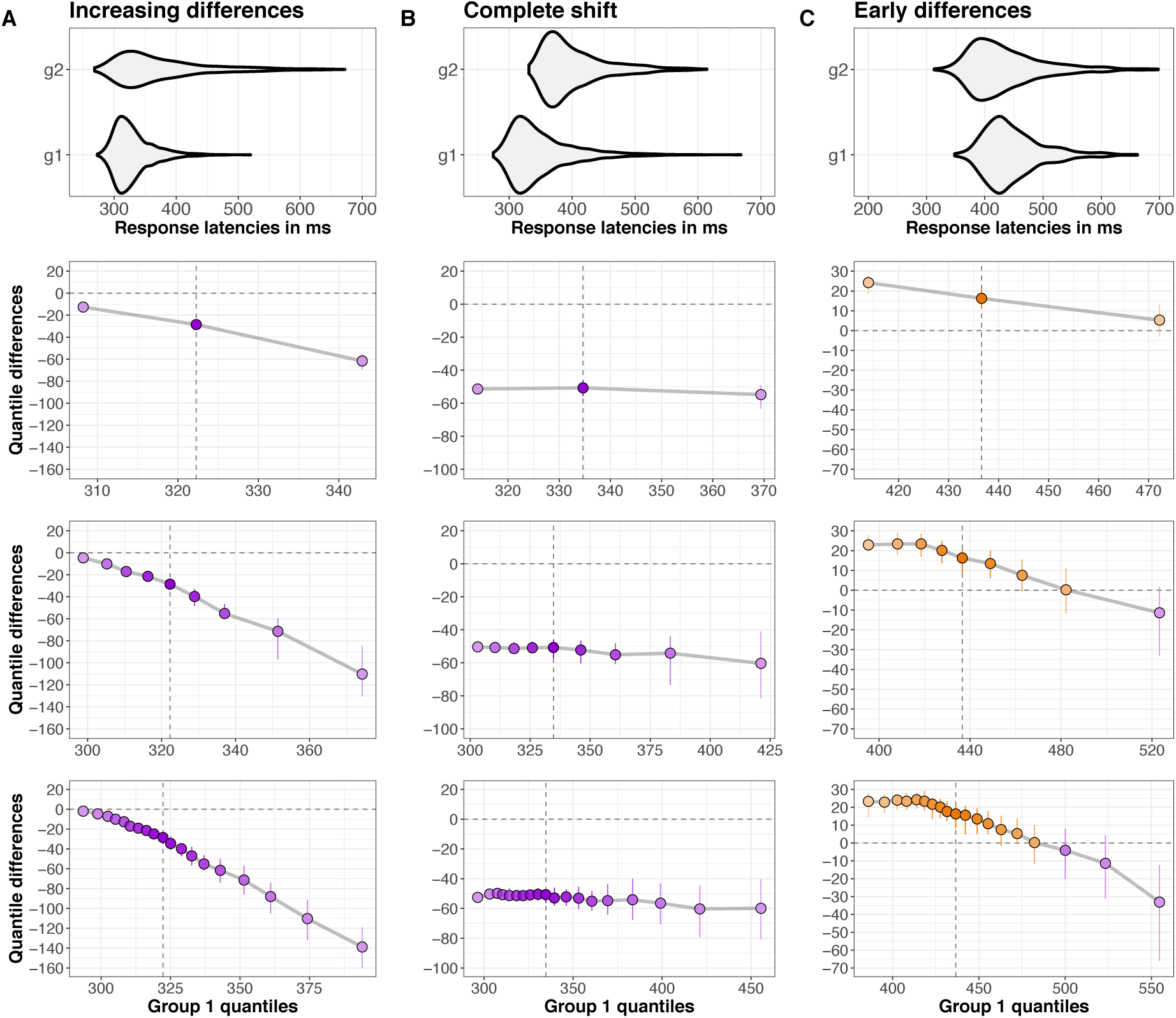
Examples of different ways in which two response time distributions could differ. **A.** Weak early differences, then increasing differences for longer latencies. **B.** Complete shift. **C.** Large early differences, then decreasing differences for longer latencies. The top row shows violinplots contrasting two distributions in the different situations. The remaining rows show shift functions with different densities applied to the same data. Row 2 estimates only the quartiles, row 3 quantifies the deciles, and row 4 quantifies quantiles 0.05 to 0.95 in steps of 0.05.

Such detailed dissociations have been reported in the literature, and provide much stronger constraints on the underlying cognitive architecture than comparisons limited to say the median reaction times across participants (Ridderinkhof *et al.*, 2005; Pratte *et al.*, 2010). A similar approach could be applied to various types of behavioural and neuronal response times and response durations.

## Perspectives on independent groups

Now that we have introduced shift functions, we need to step back to consider the different perspectives we can have when comparing two groups, starting with independent groups. So far, we have focused on *Question 1* introduced earlier: how does the typical observation in one group compares to the typical observation in the other group? *Question 2* addresses an alternative approach: what is the typical difference between any member of group 1 and any member of group 2?

Let’s look at the example in Figure 7, showing two independent samples. The scatterplots indicate large differences in spread between the two groups, and suggest larger differences in the right than the left tails of the distributions. The medians of the two groups are very similar, so the two distributions do not seem to differ in central tendency. In keeping with these observations, a t-test and a Mann-Whitney-Wilcoxon test are not significant, but a Kolmogorov-Smirnov test is.

**Figure 7.**
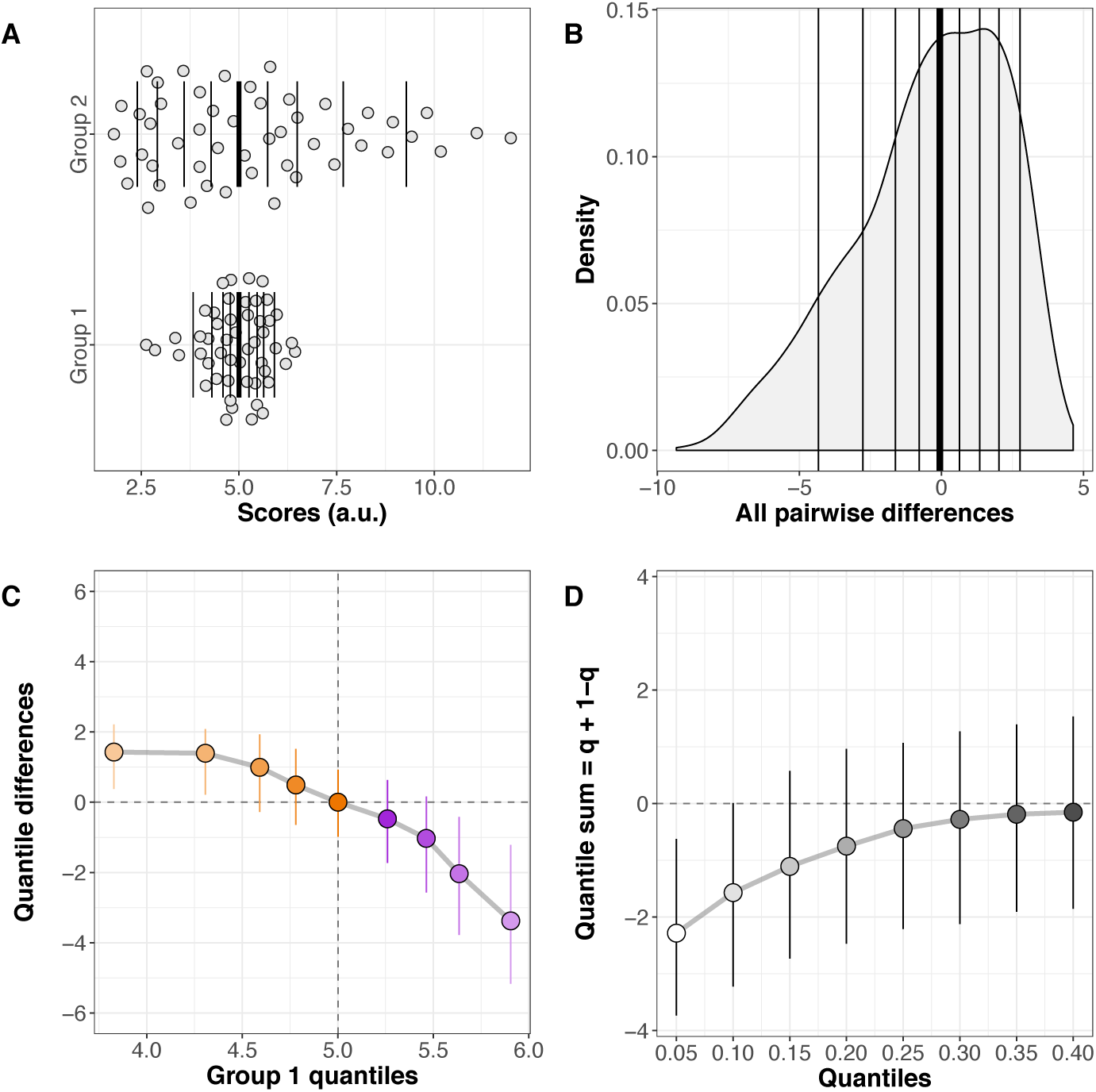
How two independent distributions differ. **A.** Stripcharts of marginal distributions. Vertical lines mark the deciles, with a thicker line for the median. **B.** Kernel density representation of the distribution of all pairwise differences between the two groups. Vertical lines mark the deciles, with a thicker line for the median. **C.** Shift function. Group 1 - group 2 is plotted along the y-axis for each decile, as a function of group 1 deciles. For each decile difference, the vertical line indicates its 95% bootstrap confidence interval. The 95% confidence intervals are controlled for multiple comparisons. **D.** Difference asymmetry function with 95% confidence intervals. The family-wise error is controlled by adjusting the critical p values using Hochberg’s method; the confidence intervals are not adjusted.

This discrepancy between tests highlights an important point: if a t-test is not significant, one cannot conclude that the two distributions do not differ. A shift function helps us understand how the two distributions differ (Figure 7C): the overall profile corresponds to two centred distributions that differ in spread; for each decile, we can estimate by how much they differ, and with what uncertainty; finally, the non-linear shift function indicates that the differences in spread are asymmetric, with larger differences in the right tails of the marginal distributions.

To address *Question 2*, we compute all the pairwise differences between members of the two groups. In this case, each group has n = 50, so we end up with 2,500 differences. Figure 7B shows a kernel density representation of these differences. What does the typical difference look like? The median of the differences is very near zero, at −0.06, with a 95% confidence interval of [−1.02, 0.75]. So, it seems on average, if we randomly select one observation from each group, they will differ very little. However, the differences can be quite substantial, and with real data we would need to put these differences in context, to understand how large they are, and their physiological interpretation. The differences are also asymmetrically distributed: negative scores extend to −10, whereas positive scores don’t even reach + 5. In other words, negative differences tend to outweigh positive differences. This asymmetry relates to our earlier observation of asymmetric differences in the shift function. If the two distributions presented in Figure 7A did not differ, the distribution of all pairwise differences should be approximately symmetric and centred about zero. Thus, the two distributions seem to differ, but in a way that is not captured by measures of central tendency.

Recently, Wilcox suggested a new approach to quantify asymmetries in difference distributions like the one in Figure 7B (Wilcox, 2012). The idea is to get a sense of the asymmetry of the difference distribution by computing a sum of quantiles = q + (1-q), for various quantiles estimated using the Harrell-Davis estimator. A percentile bootstrap technique is used to derive confidence intervals. Figure 7D shows the resulting difference asymmetry function. In this plot, 0.05 stands for the sum of quantile 0.05 + quantile 0.95; 0.10 stands for the sum of quantile 0.10 + quantile 0.90; and so on… The approach is not limited to these quantiles, so sparser or denser functions could be tested too. Figure 7D reveals negative sums of the extreme quantiles (0.05 + 0.95), and progressively smaller, converging to zero sums as we get closer to the centre of the distribution. If the distributions did not differ, the difference asymmetry function would be expected to be about flat and centred near zero. So, the q+(1-q) plot suggests that the distribution of differences is asymmetric, based on the 95% confidence intervals: the two groups seem to differ, with maximum differences in the tails. Other alpha levels can be assessed too.

## Perspectives on dependent groups

The tools covered so far have versions for dependent groups as well. Let’s consider the dataset presented in Figure 8. Panel A shows the two distributions, with relatively large differences in the right tails. To address *Question 1*, ‘How does the typical observation in condition 1 compare to the typical observation in condition 2?’, we consider the median of each condition. In condition 1 the median is 12.1; in condition 2 it is 14.8. The difference between the two medians is −2.73, with a 95% confidence interval of [−6.22, 0.88], thus suggesting a small difference between marginal distributions. To complement these descriptions, we consider the shift function for dependent groups (Wilcox & Erceg-Hurn, 2012). The shift function (Figure 6E) addresses an extension of *Question 1*, by more systematically comparing the distributions. It shows a non-uniform shift between the marginal distributions: the first three deciles do not differ significantly, the remaining deciles do, and there is an overall trend of growing differences as we progress towards the right tails of the distributions. In other words, among larger observations, observations in condition 2 tend to be higher than in condition 1.

**Figure 8.**
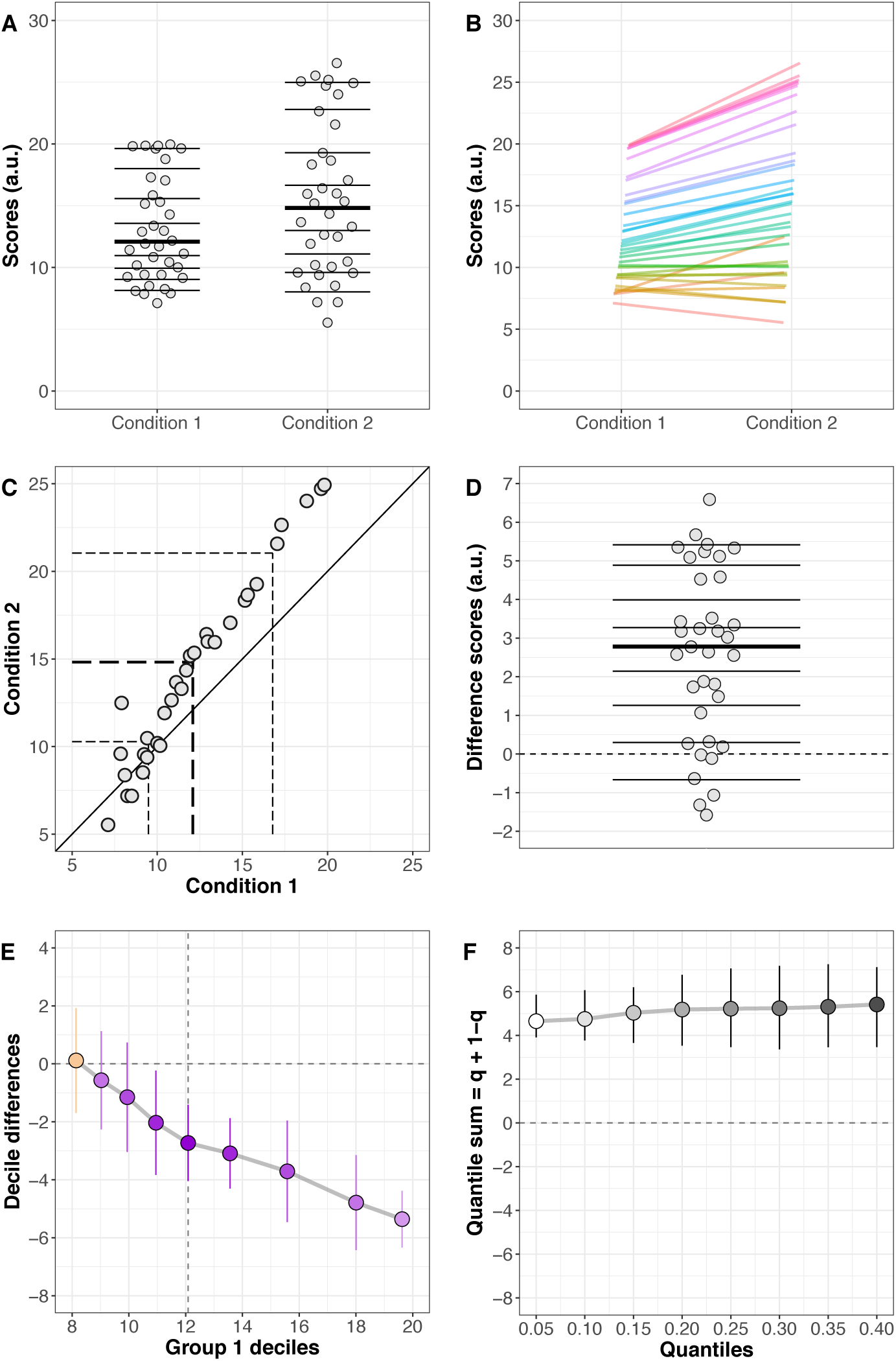
How two dependent distributions differ. **A.** Stripcharts of the two distributions. Horizontal lines mark the deciles, with a thicker line for the median. **B.** Lines joining paired observations. Scatter was introduced along the x axis to reveal overlapping observations. **C.** Scatterplot of paired observations. The diagonal black reference line of no effect has slope one and intercept zero. The dashed lines mark the quartiles of the two conditions. In panel C, it could also be useful to plot the pairwise differences as a function of condition 1 results. **D.** Stripchart of difference scores. Horizontal lines mark the deciles, with a thicker line for the median. **E.** Shift function with 95% confidence intervals. **F.** Difference asymmetry function with 95% confidence intervals.

Because we are dealing with a paired design, our investigation should not be limited to a comparison of the marginal distributions; it is also important to show how observations are linked between conditions. This association is revealed in two different ways in panels B & C. Looking at the pairing reveals a pattern otherwise hidden: for participants with weak scores in condition 1, differences tend to be small and centred about zero; beyond a certain level, with increasing scores in condition 1, the differences get progressively larger.

Panel D shows the distribution of these differences, which let us assess *Question 2*, ‘What is the typical difference for a randomly sampled participant?’. The distribution of within-participant differences is shifted up from zero, with only 6 out of 35 differences inferior to zero. Matching this observation, only the first decile is inferior to zero. The median difference is 2.78, and its 95% confidence interval is [1.74, 3.53]. To complement these descriptions of the difference distribution, we consider the difference asymmetry function for dependent groups (Wilcox & Erceg-Hurn, 2012). The difference asymmetry function extends *Question 2* about the typical difference, by considering the symmetry of the distribution of differences. In the case of a completely ineffective experimental manipulation, the distribution of differences should be approximately symmetric about zero. The associated difference asymmetry function should be flat and centred near zero. For the data at hand, Figure 8F reveals a positive and almost flat function, suggesting that the distribution of differences is almost uniformly shifted away from zero. If some participants had particularly large differences, the left part of the difference asymmetry function would be shifted up compare to the rest of the function, a non-linearity that would suggest that the differences are not symmetrically distributed – this does not seem to be the case here.

Finally, given a sufficiently large sample size, a single distribution of differences such as the one shown in Figure 8D can be quantified in more details, by including confidence intervals of the quantiles. Figure 9 illustrates such detailed representation using event-related potential onsets from 120 participants (Bieniek *et al.*, 2016). In that case, the earliest latencies are particularly interesting, so it is useful to quantify the first deciles in addition to the median.

**Figure 9.**
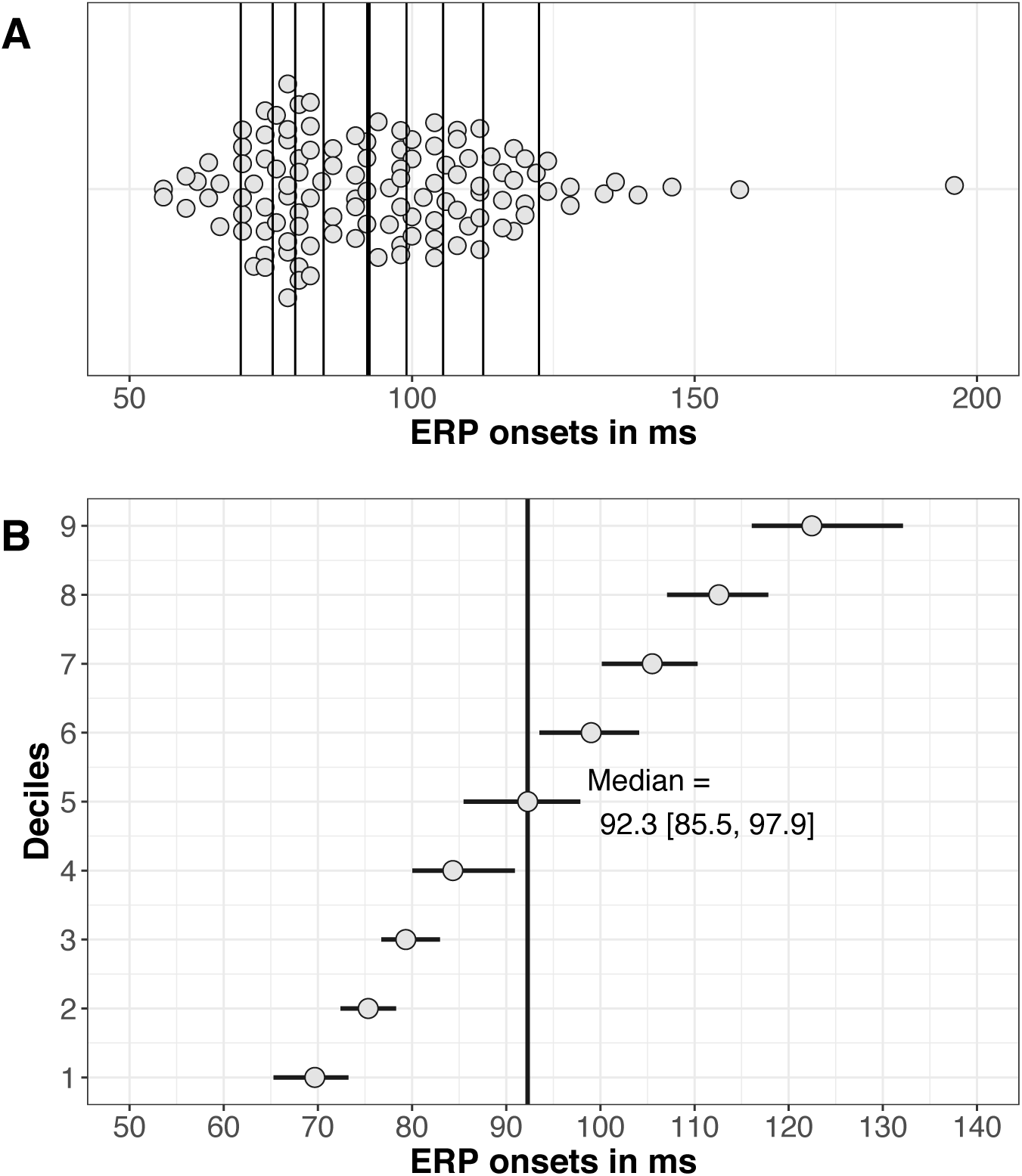
Detailed quantification of a single distribution. **A.** The scatterplot illustrates the distribution of event-related potential (ERP) onsets. Points were scattered along the y-axis to avoid overlap. Vertical lines indicate the deciles, with the median shown with a thicker line. One outlier (>200 ms) is not shown. **B.** Deciles and their 95% percentile bootstrap confidence intervals are superimposed. The vertical black line marks the median.

## Recommendations

There are various ways to illustrate and compare distributions, including how to compute a shift function and a difference asymmetry function. Therefore, the examples presented in this article should be taken as a starting point, not as a definitive answer to the experimental situations we considered. For instance, although powerful, Wilcox’s 1995 shift function technique is limited to the deciles, can only be used with alpha = 0.05, and does not work well with tied values. To circumvent these problems, Wilcox’s recently proposed a new version of the shift function that uses a straightforward percentile bootstrap without estimation of the standard error of the decile differences (Wilcox & Erceg-Hurn, 2012; Wilcox *et al.*, 2014). This new approach allows tied values, can be applied to any quantile and can have more power when looking at extreme quantiles (q<=0.1, or q>=0.9). This version of the shift function gives the opportunity to quantify the effects at different resolutions, to create sparser or denser shift functions, as demonstrated in Figure 6. The choice of resolution depends on the application, the precision of the hypotheses, and the sample size. For dependent variables, at least 30 observations are recommended to compare the 0.1 or 0.9 quantiles (Wilcox & Erceg-Hurn, 2012). To compare the quartiles, 20 observations appear to be sufficient. The same recommendations hold for independent variables; in addition, to compare the .95 quantiles, at least 50 observations per group should be used (Wilcox *et al.*, 2014). For the difference asymmetry function, if sample sizes are equal, it seems that n = 10 is sufficient to assess quantiles 0.2 and above. To estimate lower quantiles, n should be at least 20 in each group (Wilcox, 2012). Such large numbers of observations might seem daunting in certain fields, but there is simply no way around this fundamental limitation: the more precise and detailed our inferences, the more observations we need.

## Conclusion

The techniques presented here provide a very useful perspective on group differences, by combining detailed illustrations and quantifications of the effects. The different techniques address different questions, so which technique to use depends on what is the most interesting question in a particular experimental context. This choice should be guided by experience: to get a good sense of the behaviour of these techniques requires practice with both real and simulated data. By following that path, the community will soon realise that classic approaches such as t-tests on means combined with bar graphs are far too limited, and richer information can be captured in datasets, which in turn can lead to better theories and understanding of the brain.

One might think that such detailed analyses will increase false positives, and risk to focus on trivial effects. However, the tools presented here control for multiple comparisons, thus limiting false positives. Nevertheless, applying multiple tests to the same dataset, such as a t-test, a Kolmogorov-Smirnov test, a shift function, and difference asymmetry function, will inevitably increase false positives. There is a simple safeguard against these problems: replication. Drawing inspiration from genetic studies, we should consider two samples, one for discovery, one for replication. The tools described in this article are particularly useful to explore distributions in a discovery sample. Effects of interest can then be tested in a replication sample. Our approach has also the advantage of taking the focus away from binary outcomes (significant vs. non-significant), towards robust effect sizes and the quantification of exactly how distributions differ.

## Acknowledgements

We thank Tracey Weissgerber and Richard Morey for their very constructive and detailed reviews of previous versions of this article. Readers can see the original version of the article on figshare to appreciate how much the review process improved our paper: https://doi.org/10.6084/m9.figshare.4055970.v1

## References

Allen, E.A., Erhardt, E.B. & Calhoun, V.D. (2012) Data visualization in the neurosciences: overcoming the curse of dimensionality. Neuron, 74, 603–608.

Anscombe, F.J. (1973) Graphs in Statistical Analysis. Am Stat, 27, 17–21.

Bieniek, M.M., Bennett, P.J., Sekuler, A.B. & Rousselet, G.A. (2016) A robust and representative lower bound on object processing speed in humans. The European journal of neuroscience, 44, 1804–1814.

Bjerkedal, T. (1960) Acquisition of resistance in guinea pigs infected with different doses of virulent tubercle bacilli. Am J Hyg, 72, 130–148.

Button, K.S., Ioannidis, J.P., Mokrysz, C., Nosek, B.A., Flint, J., Robinson, E.S. & Munafo, M.R. (2013) Power failure: why small sample size undermines the reliability of neuroscience. Nature reviews. Neuroscience, 14, 365–376.

Chanauria, N., Bharmauria, V., Bachatene, L., Cattan, S., Rouat, J. & Molotchnikoff, S. (2016) Comparative effects of adaptation on layers II-III and V-VI neurons in cat V1. The European journal of neuroscience, 44, 3094–3104.

Clarke, E. & Sherrill-Mix, S. (2016) ggbeeswarm: Categorical Scatter (Violin Point) Plots. R package version 0.5.3. https://cran.r-project.org/package=ggbeeswarm

Colquhoun, D. (2014) An investigation of the false discovery rate and the misinterpretation of p-values. R Soc Open Sci, 1, 140216.

Cook, D., Lee, E.K. & Majumder, M. (2016) Data Visualization and Statistical Graphics in Big Data Analysis. Annu Rev Stat Appl, 3, 133–159.

De Jong, R., Liang, C.C. & Lauber, E. (1994) Conditional and Unconditional Automaticity - a Dual-Process Model of Effects of Spatial Stimulus - Response Correspondence. J Exp Psychol Human, 20, 731–750.

Doksum, K. (1974) Empirical Probability Plots and Statistical Inference for Nonlinear Models in the two-Sample Case. Annals of Statistics, 2, 267–277.

Doksum, K.A. (1977) Some graphical methods in statistics. A review and some extensions. Statistica Neerlandica, 31, 53–68.

Doksum, K.A. & Sievers, G.L. (1976) Plotting with Confidence - Graphical Comparisons of 2 Populations. Biometrika, 63, 421–434.

Duke, S.P., Bancken, F., Crowe, B., Soukup, M., Botsis, T. & Forshee, R. (2015) Seeing is believing: good graphic design principles for medical research. Stat Med, 34, 3040–3059.

Erceg-Hurn, D.M. & Mirosevich, V.M. (2008) Modern robust statistical methods: an easy way to maximize the accuracy and power of your research. Am Psychol, 63, 591–601.

Forstmeier, W., Wagenmakers, E.J. & Parker, T.H. (2016) Detecting and avoiding likely false-positive findings - a practical guide. Biol Rev Camb Philos Soc.

Handcock, M.S. & Morris, M. (1998) Relative distribution methods. Sociol Methodol, 28, 53–97.

Harrell, F.E. & Davis, C.E. (1982) A new distribution-free quantile estimator. Biometrika, 69, 635–640.

Ince, R.A.A., Giordano, B.L., Kayser, C., Rousselet, G.A., Gross, J. & Schyns, P.G. (2016) A statistical framework for neuroimaging data analysis based on mutual information estimated via a Gaussian copula. bioRxiv.

Massidda, D. (2013) retimes: Reaction Time Analysis. R package version 0.1–2. https://cran.r-project.org/package=retimes

Munafò, M.R., Nosek, B.A., Bishop, D.V.M., Button, K.S., Chambers, C.D., Percie du Sert, N., Simonsohn, U., Wagenmakers, E.-J., Ware, J.J. & Ioannidis, J.P.A. (2017) A manifesto for reproducible science. Nature Human Behaviour, 1, 0021.

Pernet, C., Andersson, J., Paulesu, E. & Demonet, J.F. (2009a) When all hypotheses are right: a multifocal account of dyslexia. Hum Brain Mapp, 30, 2278–2292.

Pernet, C.R., Poline, J.B., Demonet, J.F. & Rousselet, G.A. (2009b) Brain classification reveals the right cerebellum as the best biomarker of dyslexia. BMC Neurosci, 10, http:--http://www.biomedcentral.com-1471-2202-1410-1467-/doi:1410.1186-1471-2202-1410-1467.

Poldrack, R.A., Baker, C.I., Durnez, J., Gorgolewski, K.J., Matthews, P.M., Munafo, M.R., Nichols, T.E., Poline, J.B., Vul, E. & Yarkoni, T. (2017) Scanning the horizon: towards transparent and reproducible neuroimaging research. Nature reviews. Neuroscience, 18, 115–126.

Pratte, M.S., Rouder, J.N., Morey, R.D. & Feng, C.N. (2010) Exploring the differences in distributional properties between Stroop and Simon effects using delta plots. Atten Percept Psycho, 72, 2013–2025.

R Core Team 2016) R: A Language and Environment for Statistical Computing. https://www.r-project.org/

Ridderinkhof, K.R., Scheres, A., Oosterlaan, J. & Sergeant, J.A. (2005) Delta plots in the study of individual differences: New tools reveal response inhibition deficits in AD/HD that are eliminated by methylphenidate treatment. J Abnorm Psychol, 114, 197–215.

Rousselet, G., Pernet, C. & Wilcox, R. (2016a) Modern graphical methods to compare two groups of observations. figshare. https://dx.doi.org/10.6084/m9.figshare.4055970

Rousselet, G.A., Foxe, J.J. & Bolam, J.P. (2016b) A few simple steps to improve the description of group results in neuroscience. The European journal of neuroscience, 44, 2647–2651.

Rousselet, G.A. & Wilcox, R.R. (2016) rogme: Robust Graphical Methods For Group Comparisons. R package version 0.1.0.9000. https://github.com/GRousselet/rogme

Speckman, P.L., Rouder, J.N., Morey, R.D. & Pratte, M.S. (2008) Delta plots and coherent distribution ordering. Am Stat, 62, 262–266.

Talebi, V. & Baker, C.L., Jr. (2016) Categorically distinct types of receptive fields in early visual cortex. J Neurophysiol, 115, 2556–2576.

Wagenmakers, E.J., Wetzels, R., Borsboom, D., van der Maas, H.L. & Kievit, R.A. (2012) An Agenda for Purely Confirmatory Research. Perspect Psychol Sci, 7, 632–638.

Weissgerber, T.L., Milic, N.M., Winham, S.J. & Garovic, V.D. (2015) Beyond bar and line graphs: time for a new data presentation paradigm. PLoS Biol, 13, e1002128.

Wickham, H. (2016) ggplot2: Elegant Graphics for Data Analysis. Springer International Publishing.

Wilcox, R.R. (1995) Comparing Two Independent Groups Via Multiple Quantiles. Journal of the Royal Statistical Society. Series D (The Statistician), 44, 91–99.

Wilcox, R.R. (2001) Modern insights about Pearson’s correlation and least squares regression. Int J Select Assess, 9, 195–205.

Wilcox, R.R. (2006) Graphical methods for assessing effect size: Some alternatives to Cohen’s d. Journal of Experimental Education, 74, 353–367.

Wilcox, R.R. (2009) Basic statistics: understanding conventional methods and modern insights. Oxford University Press, New York; Oxford.

Wilcox, R.R. (2012) Comparing Two Independent Groups Via a Quantile Generalization of the Wilcoxon-Mann-Whitney Test. Journal of Modern Applied Statistical Methods, 11, 296–302.

Wilcox, R.R. (2017) Introduction to Robust Estimation and Hypothesis Testing. Academic Press, 4th edition.

Wilcox, R.R. & Erceg-Hurn, D.M. (2012) Comparing two dependent groups via quantiles. J Appl Stat, 39, 2655–2664.

Wilcox, R.R., Erceg-Hurn, D.M., Clark, F. & Carlson, M. (2014) Comparing two independent groups via the lower and upper quantiles. J Stat Comput Sim, 84, 1543–1551.

Wilcox, R.R. & Keselman, H.J. (2003) Modern Robust Data Analysis Methods: Measures of Central Tendency. Psychological Methods, 8, 254–274.

Wilcox, R.R. & Muska, J. (2010) Measuring effect size: A non-parametric analogue of omega(2). The British journal of mathematical and statistical psychology, 52, 93–110.

Wilke, C.O. (2016) cowplot: Streamlined Plot Theme and Plot Annotations for ‘ggplot2’. R package version 0.6.2. https://cran.r-project.org/package=cowplot

